# Bayesian Gaussian Process Latent Variable Models for pseudotime inference in single-cell RNA-seq data

**DOI:** 10.1101/026872

**Authors:** Kieran Campbell, Christopher Yau

## Abstract

Single-cell genomics has revolutionised modern biology while requiring the development of advanced computational and statistical methods. Advances have been made in uncovering gene expression heterogeneity, discovering new cell types and novel identification of genes and transcription factors involved in cellular processes. One such approach to the analysis is to construct pseudotime orderings of cells as they progress through a particular biological process, such as cell-cycle or differentiation. These methods assign a score - known as the pseudotime - to each cell as a surrogate measure of progression. However, all published methods to date are purely algorithmic and lack any way to give uncertainty to the pseudotime assigned to a cell. Here we present a method that combines Gaussian Process Latent Variable Models (GP-LVM) with a recently published electroGP prior to perform Bayesian inference on the pseudotimes. We go on to show that the posterior variability in these pseudotimes leads to nontrivial uncertainty in the pseudo-temporal ordering of the cells and that pseudotimes should not be thought of as point estimates.

## 1 Introduction

Single-cell RNA-seq (scRNA-seq) has emerged as a powerful method for the quantification of gene and transcript abundance in individual cells. In only a few years it has uncovered exciting new biology including the identification of novel cell types [1], hidden heterogeneity in gene expression [2] and regulatory networks operating at the single-cell level [3]. It has particular advantages over bulk RNA sequencing such as the ability to identify rare cell types and cell-to-cell variability [4, 5], which are typically hidden in bulk analyses.

Despite being simultaneously sequenced individual cells may be of variable progression through a variety of cellular processes due to heterogeneous responses to stimuli and inherent transcriptomic variability. This has consequently lead to the idea of pseudotime as an artificial measure of a cell’s progression through a process such as differentiation or apoptosis [6]. The statistical problem is to assign a pseudotime label between 0 and 1 to each observation where values near 0 indicates that the cell is in a state near the start of the biological process and values near 1 denote cells that are toward the end of the process.

Early attempts and pseudotime ordering algorithms include *Monocle*[6], which uses Independent Component Analysis (ICA) with Minimum Spanning Trees (MST) applied to scRNA-seq data, *Wanderlust* [7], which uses ensembles of *k*-nearest-neighbour graphs applied to mass spectrometry data and *embeddr* [8], which uses Laplacian Eigenmaps and principal curves to identify pseudotime trajectory using nonparametric curve fitting. However, a common feature of all these algorithms is that they provide point estimates of the pseudotime for each cell and are unable to quantify the uncertainty in each estimate^1^. As a result, the differential expression of particular genes over pseudotime may simply be an artefact of that ordering and disappear when the uncertainty is taken into account.

Here we present an innovative approach that uses full Bayesian inference for Gaussian Process Latent Variable Models [10] combined with a repulsive prior [11] to infer posterior distributions of pseudotimes. First, a reduced dimension representation of the cells from a dimensionality reduction algorithm such as principal components analysis (PCA) or Laplacian Eigenmaps [12] is created. Subsequently, a probabilistic curve is fitted through the cells that jointly fits posterior distributions of pseudotimes over all cells. As a result the uncertainty in each pseudotime assignment can be accurately assessed as well as the extent to which the ordering of any two cells is robust against the inherent noise. Our method does not require cell capture times as a prerequisite. An implementation of our inference method is available in the Julia programming language at <u>http://github.com/kieranrcampbell/gpseudotime</u>.

## 2 Gaussian Processes for pseudo-time assignment

### 2.1 Gaussian Process Latent Variable Models

A real-valued stochastic process {*μ*_*t*_, *t* ∈ *T*}, where *T* is an index set, is a Gaussian Process (GP), if all its finite dimensional marginal distributions are multivariate Gaussian distributions. That is, for any given distinct values *t*_1_, *…, t*_*n*_, the random vector ***μ*** = (*μ*_*t*_1__, *…, μ*_*t*_*n*__) *∼ N* (**m**, **K**) where **m** *≡* 𝔼[***μ*** ] and **K** *≡* cov(***μ***, ***μ***).

Gaussian Processes allow us to define prior probability distributions over real-valued functions in Bayesian nonparametric modelling. A popular model exploiting this property is the Gaussian Process Latent Variable Model (GPLVM) [13]. GPLVMs are a family of non-parametric methods that define a distribution over functions *μ* linking a latent variable *t* to an output variable *x*, for example, through the relationship *x* = *μ* (*t*) + *ϵ* where *ϵ ∼ 𝒩* (0, *σ*^2^). If the prior distribution over the latent function *μ* is given by a Gaussian Process then, given a set of *N* observations **x** = *{x*_*t*_1__, *…, x*_*t*_N__} and corresponding latent points **t** = *{t*_1_, *…, t*_*N*_}, the marginal distribution *p*(**x**|**t**) is multivariate Gaussian with mean vector **m**(**t**) and covariance matrix given by **K** + *σ*^2^***I***_*N*_ where *K*_*ij*_ = *κ*(*t*_*i*_, *t*_*j*_) for a kernel function *κ* and ***I***_*N*_ is an *N × N* identity matrix [10]. The specification of the kernel function *κ* allows us to control the favourable features of the latent functions, e.g. smoothness.

### 2.2 Model Specification

In our problem, the data consists of *N P*-dimensional observation vectors 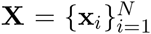 which are assumed to be conditionally independent given the latent, unobserved pseudo-times **t** = {*t*_1_, *… t_n_*}, *t_i_* ∈ (0, 1], a mean function ***μ*** (*t*) and a observation covariance matrix **∑**. The data we begin with are ‘features’ measured across *N* cells that we wish to order. These features can either be genes of particular interest or co-ordinates of some previously applied manifold learning algorithm, such as Laplacian Eigenmaps [12]. Each dimension of the mean function *μ*_*j*_ is given an independent Gaussian Process prior with covariance function *K*^(*j*)^.

Our model is described succinctly in the following hierarchical representation:

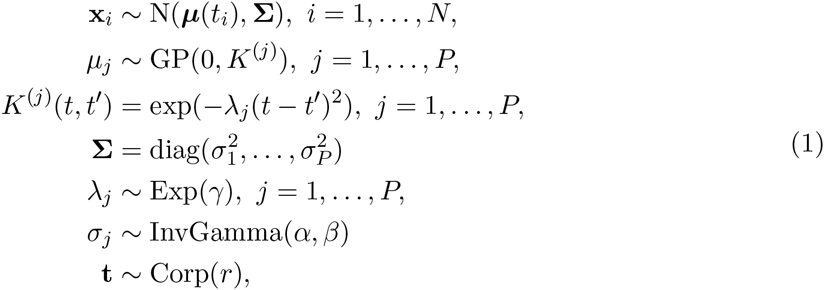

where in the last line we define a repulsive prior on the pseudo-times **t** based on the Coulomb repulsive process [11].

The Coulomb repulsive process models repulsions between adjacent points using a process inspired by physical models of electrostatic potentials. In our model, this has the effect of preferentially favouring pseudo-time configurations that ‘fill out’ the interval (0, 1]. For pseudotimes *t*_1_, *…, t*_*N*_ the prior is given by

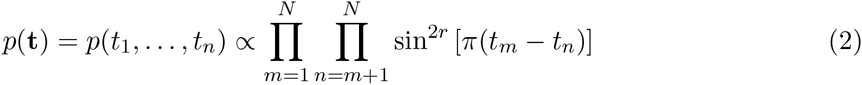

where *r* is the repulsion parameter. Under this prior, as *|t*_*i*_ − *t*_*j*_| gets smaller, the probability decreases and there is zero probability that two pseudo-times will coincide. [11] demonstrated that embedding this process within a GPLVM gave superior results to standard alternatives such as uniform *t*_*i*_ *∼ U* (0, 1) or normal priors *t*_*i*_ *∼ N* (0, 1) by avoiding identifiability issues due to scale- and translational-invariance under these standard priors.

The likelihood of **X** given the latent pseudotimes **t** is conditionally independent across features [10] so we can write it as

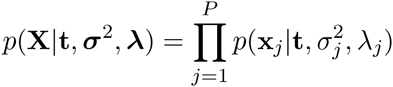

where

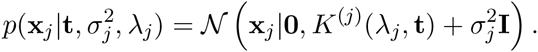

Therefore, we can write the joint posterior as

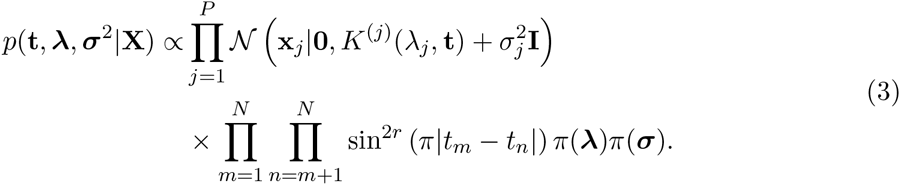

Note that for maximum likelihood inference **t** will never leave its initial ordering as the prior goes to zero probability when *t*_*m*_ = *t*_*n*_.

Interestingly, the parameter ***λ*** has an intuitive interpretation in the context of curve fitting. In a one-dimensional Gaussian Process regression setting, *λ* corresponds to the ‘horizontal’ length scale over which the function varies. Therefore, in the two-dimensional plane |***λ***| loosely corresponds to arc-length, with larger |***λ***| generating longer curves. We effectively regularize ***λ*** by placing an exponential prior on it, with larger *γ* corresponding to shorter curves passing through the manifold.

### 2.3 Statistical Inference

Statistical inference for GPLVMs is typically performed using approximate maximum *a posteriori* [11] or variational methods [10] but Markov Chain Monte Carlo (MCMC) approaches are also possible [14]. As our primary objective is to characterise full posterior uncertainty measures of pseudo-time, MCMC-based inference was a necessity.

We therefore adopted used a random-walk Metropolis-Hastings (MH) with normal proposals for the unknown parameters of the model (***t***, ***λ***, ***σ***^2^):

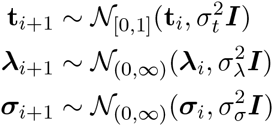

where 𝒩_[*a,b*]_ is the multivariate Gaussian distribution truncated on [*a, b*]. The marginal properties of the Gaussian Process meant that the latent functions *μ* could be integrated out as therefore we did not need to impute this infinite-dimensional quantity.

One difficulty with the proposed model is the extreme multi-modal nature of the posterior. Many local maxima exist in the prior alone due to the *N* (*N* − 1)/2 values for which it vanishes whenever *t*_*i*_ = *t*_*j*_. As such, setting initial values to avoid becoming stuck in local maxima is challenging^2^. However, we can exploit the *repulsive* nature of the prior, by initialising 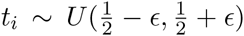 for arbitrarily small *ϵ*. As the prior heavily discourages pseudo-times that are in close proximity, the prior pushes the pseudotimes apart like charged particles being repelled while the likelihood biases this in an order that is consistent with the data reflecting the *a posteriori* more likely pseudotime orderings. We call this a “Big Bang” initialisation using an analogy to the famous cosmological phenomenon. Note that in practice it is really the variance 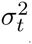 in the proposal distribution for **t** that provides the spread of points at the first iteration rather than the value of *ϵ* itself, so any sensible *ϵ < σ*_*t*_ will work.

## 3 Results

We applied our method to two datasets: (i) a synthetic dataset generated from the model, and (ii) a Laplacian Eigenmaps representation of the *Monocle* [6] dataset of differentiating myoblasts.

### 3.1 Synthetic Data

We generated data from the Gaussian Process described in Equation 1. Specifically, we set ***λ*** = [1, 2], ***σ*** = [2 *×* 10^−3^, 2 *×* 10^−2^] and sampled *n* = 100 pseudotimes from U(0, 1). We performed MH inference as described in Section 2.3 using 2 *×* 10^5^ iterations thinned by 100. We set *ϵ* = 10^−6^, *σ*_*t*_ = 9 *×* 10^−3^, *σ*_*λ*_ = 5 *×* 10^−1^ and *σ*_*σ*_ = 5 *×* 10^−3^. The hyper-parameters were set to *r* = 10^−3^, *α* = *β* = *γ* = 1. The results of inference on the synthetic data can be seen in Figure 1. Pseudotime estimates clearly converge to stable values that are close to the ‘true’ values and are almost always within the 95% highest probability density (HPD) credible interval.

**Figure 1:**
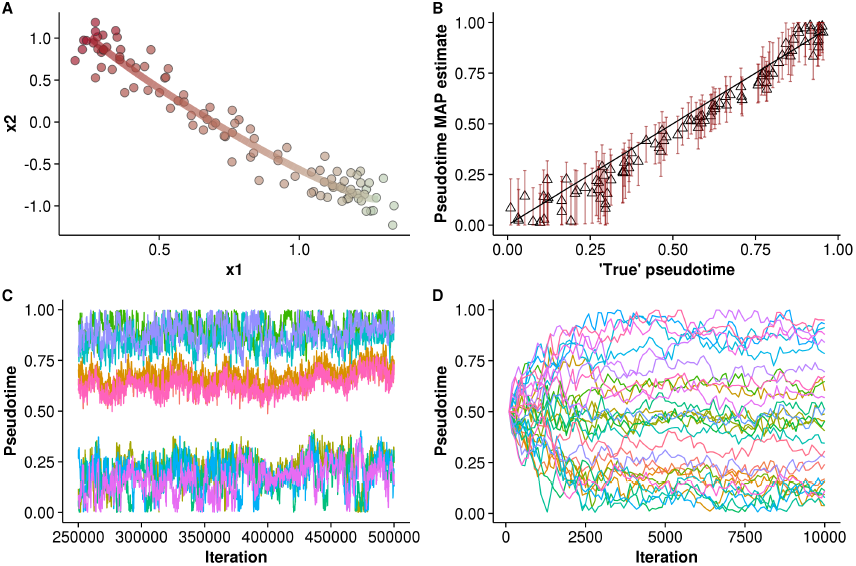
Pseudotime inference on synthetic data. **A.** The synthetic data points along with the MAP estimate of the mean function *μ* (*t*). Points are coloured by ‘true’ pseudotime while the curve is coloured by the MAP estimate. **B.** ‘True’ pseudotimes plotted against the MAP pseudotime estimates. Error bars report the 95% HPD credible interval; the solid line corresponds to *y* = *x*. **C.** MCMC traces for ten randomly chosen cells after the burn-in period. **D.** The ‘big bang’ intialisation for thirty randomly chosen cells up to 10^4^ iterations.

### 3.2 Single-cell RNA-seq dataset

We next applied our method to a dataset of differentiating myoblasts [6]. A Laplacian Eigenmaps reduced-dimensionality representation of the data was used, as described at 

~~~
https://github.com/kieranrcampbell/embeddr/blob/master/vignettes/vignette.Rmd
~~~

. Four outlier cells were discarded and the points were centre scaled to have mean 0 and standard deviation 1 in each dimension. We performed MH inference as described in Section 2.3 using 5 *×* 10^5^ iterations thinned by 500. We set *ϵ* = 10^−6^, *σ*_*t*_ = 6.5 *×* 10^−3^, *σ*_*λ*_ = 9 *×* 10^−13^ and *σ*_*σ*_ = 8 *×* 10^−3^. The hyper-parameters were set to *r* = 10, *α* = *β* = 1, *γ* = 100.

The results of the inference can be seen in Figure 2. Clearly the GP-LVM curve traces through the centre of the manifold in a manner similar to the principal curve fit obtained using the 

~~~
embeddr
~~~

 [8] (Figure 2A). The MAP pseudotime values are very similar to the principal curve values with almost all falling within the 95% CI (Figure 2B).^3^

**Figure 2:**
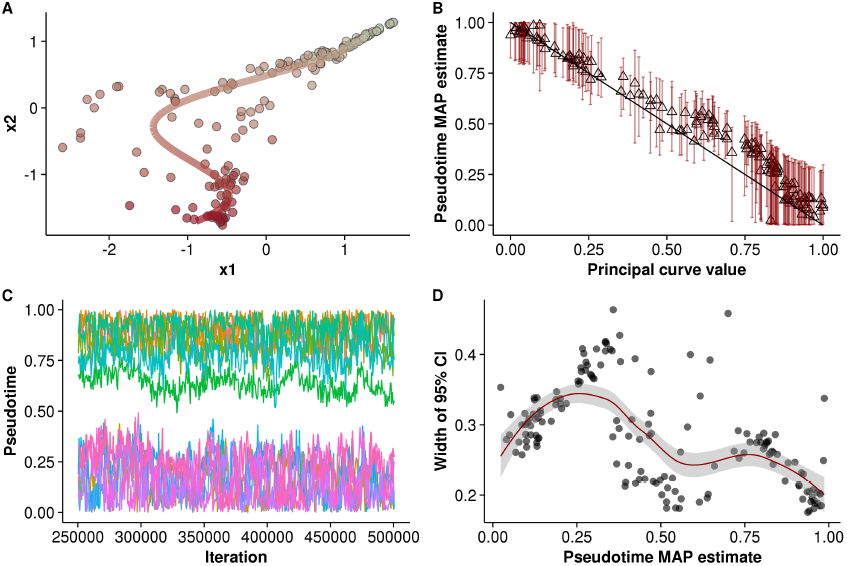
Pseudotime inference on scRNA-seq data of differentiating myoblasts. **A.** The Laplacian Eigenmaps representation of the dataset, with each point coloured by the MAP pseudotime estimate, along with the posterior mean curve of the GPLVM. **B.** The MAP pseudotime estimates compared to the principal curve fit, along with the 95% HPD credible interval. **C.** The thinned MCMC traces after burn-in for 12 randomly chosen cells. **D.** The width of the 95% HPD CI as a function of pseudotime.

The variation in pseudotime estimate can be seen in Figure 2C/D. The 95% CI for some cells is as large as 0.5 - half the overall pseudotime window. This suggest that point estimation of pseudotimes could severely underestimate the potential variability in the estimates (which in the point estimate case is assumed to be negligible). Our model predicts uncertainty in such assignments can extend to almost half of the entire biological process of interest. Visually, this is reasonable since the pseudotime assignment depends on the relative positioning along a curve drawn through the data points. There are clearly a range of possible curves that could be compatible with this data and integrating out the latent function allows us to characterise this uncertainty.

An alternative way to summarise the variation is using a non-parametric measure of correlation between subsequent (thinned) MCMC pseudotime traces. We computed the mean Kendall-Tau correlation - a non-parametric measure of rank correlation - along the entire chain and found it to be 0.84. In other words, the ordering of cells consistently changes meaning the idea of a well defined ordering of cells is meaningless.

## 4 Discussion

We have presented a novel probabilistic method for inferring pseudotimes from single-cell RNA-seq data by applying Bayesian Gaussian Process Latent Variable modelling. We have shown that the use of the Coulomb repulsive prior is appropriate to provide a well defined posterior over pseudotimes. Furthermore, the use of this prior combined with the MCMC sampling can decide the ordering of points using no prior knowledge, as opposed to the method suggested in [11] where the structure of the prior combined with maximum likelihood inference means the points will never leave their initial ordering. Finally, because our method works in a reduced-dimension space an immediate visual check of plotting the MAP mean curve exists to ensure the inference method is not stuck in a local maximum.

By applying our method to both synthetic and real data we have shown that a pseudotime ordering of cells can be recovered and the inherent uncertainty in it characterised. We also uncovered a surprisingly large posterior uncertainty in pseudotimes and variability in the cell ordering, suggesting the ideas of ‘fixed’ pseudotimes should be reconsidered.

Previous approaches to pseudotime assignment have given point estimates of temporal ordering [6–8] but we have been able to infer the uncertainty associated with the pseudotime assignment to each cell with our model. Recently [9] have also used Bayesian Gaussian Process Latent Variable Models for pseudo-time assignment allowing uncertainty to be quantified. However, their approach requires the measurement of cell capture times upon which prior distributions can be centred solving identifiability issues by providing a physical calibrated temporal scaling. This information is often not available in single cell experiments. Our approach is more general providing support for data sets where actual temporal information cannot be attained. Prior capture time information can be included in our model, this would modifying the repulsive prior to be conditional on cells which have temporal information.

In future extensions of our work we will consider closer integration with the initial dimensionality reduction problem. Our simulations have assumed that the high-dimensional gene expression measurements have already been preprocessed and reduced to a low-dimensional representation (in our case using Laplacian Eigenmaps). The GPLVM framework provides a natural extension to avoid the need for this preprocessing step but inference could be challenging in the Monte Carlo sampling framework we desire. We would also like to consider improved Monte Carlo inference approaches as early experiments using standard MCMC techniques, such as parallel tempering, have yielded no significant sampling efficiency due to the extreme multimodal nature of the Coulomb repulsion prior. We are developing novel sampling techniques to address the unique properties of this prior but will also consider alternative repulsive processes. Finally, we are also examining downstream applications of our techniques for quantifying temporal gene expression behaviour.

## 5 Acknowledgements

We thank Michalis Titsias for comments and advice regarding the manuscript. K.C. is supported by the MRC. C.Y. is supported by a UK Medical Research Council New Investigator Research Grant (Ref. No. MR/L001411/1), the Wellcome Trust Core Award Grant Number 090532/Z/09/Z, the John Fell Oxford University Press (OUP) Research Fund and the Li Ka Shing Foundation via a Oxford-Stanford Big Data in Human Health Seed Grant.

## Footnotes

1 In theory a bootstrap measure of uncertainty could be assigned, though this is computationally expensive.

2 The local maxima of the posterior generally have geometrically intuitive interpretations such as the posterior mean curve folding back on itself one or more times.

3 Although it looks like the pseudotimes are reversed, since they are ‘pseudo’ by nature they’re entirely equivalent up to a parity transformation.

